# Conservation and divergence of poly(A) tail regulation during the mammalian oocyte-to-embryo transition

**DOI:** 10.1101/2021.08.29.458065

**Authors:** Yusheng Liu, Junxue Jin, Yiwei Zhang, Le-Yun Wang, Chuanxin Zhang, Zhenzhen Hou, Wei Li, Zhonghua Liu, Falong Lu, Jiaqiang Wang

## Abstract

Poly(A) tail length and non-A residues are vital for oocyte-to-embryo transition (OET) in mice and humans^1–5^. However, the role of poly(A) tail length and non-A residues during OET in other commonly used mammalian animal models for human diseases remains unexplored. In addition, the degree of conservation in maternal mRNA poly(A) tail dynamics during OET across different mammal species is unknown. Here, we conduct a comparative analysis of the poly(A) tails during OET across four species: mice, rats, pigs, and humans. Dynamics during OET found to be conserved across all four species include: maternal mRNA deadenylation during oocyte maturation and re-polyadenylation after fertilization; a fall-rise trend in poly(A) tail length distribution; a rise-fall trend in the ratio of poly(A) tails with non-A residues; higher abundance of non-A residues in poly(A) tails of maternal mRNA than in zygotic genome activation (ZGA) mRNA; maternal mRNA with U residues degrades faster than those without U residues at the stage when ZGA takes place. While in mice and rats maternal mRNA deadenylation is impaired in parthenogenetic embryos and ZGA inhibition leads to blocked maternal mRNA deadenylation in mice and humans. In contrast, the length of consecutive U residues and the duration time of U residues in poly(A) tail diverges across the four species. Together, these findings reveal that the poly(A) tail mediated maternal mRNA post-transcriptional regulation is highly conserved in mammals with unique divergences in the length and life-span of U residues, providing new insights for the further understanding of OET across different mammals.

## INTRODUCTION

The poly(A) tails found in most eukaryotic mRNAs and long non-coding RNAs (lncRNAs) are known to be essential structural components of mRNA as they play a key role in mRNA stability and translation^6–13^, and are involved in diverse biological processes including gametogenesis^14–18^, tumor metastasis^19^, innate immunity^20^, synaptic plasticity^21, 22^, and long-term memory^23^. Poly(A) tail dynamics include length changes via deadenylation and cytoplasmic polyadenylation, and the abundance of non-A residues in poly(A) tails (non-A residues for short hereafter) at 3′-end, internal, and 5′-end. The 3′-end U residues promote rapid mRNA decay^14–16, 24^, while 3′-end G residues protect mRNA from rapid deadenylation^25^. Recent technology breakthroughs have revealed widespread poly(A) tail internal non-A residues in human, mouse, and *C. elegans*^1, 26^, of which the function and synthesis mechanisms have recently started to be studied in mice and humans^1–5^.

The oocyte-to-embryo transition (OET) in mammals lays the foundation for successful reproduction^27–29^. Because transcription is silent prior to zygotic genome activation (ZGA)^13, 30^, the events during OET are controlled by the maternal RNAs, which are tightly regulated by post-transcriptional regulation mechanisms, such as dynamics of poly(A) tail length^1, 9, 29, 31, 32^. Deleting maternal *Btg4*, which encodes the adaptor of CCR4-NOT deadenylase, leads to impaired deadenylation of maternal RNAs and embryo developmental arrest during cleavage^31–33^. In addition, the U residues catalyzed by TUT4/7 and G residues catalyzed by TENT4A/B are required for mammalian OET^3, 5^.

Rats and pigs are more similar to humans in physiology and anatomy than mice and thus are considered to be excellent models for human diseases and cell therapies^34, 35^. However, the poly(A) tail dynamics during OET in pigs and rats are completely unknown and a comparative understanding of maternal mRNA poly(A) tail regulation across mammalian species is lacking. In this study, we obtained a complete picture for the first time of poly(A) tail dynamics during OET in pigs and rats, and in combination with mouse and human data^2–5^, we discovered conservation and divergences in the dynamics of poly(A) tail length and non-A residues during OET in mice, rats, pigs, and humans, which sheds new lights on our understanding of OET across different mammals.

## RESULTS

### Poly(A) tail length dynamics during pig OET

In order to explore mRNA poly(A) tail dynamics during pig OET, we applied two complementary methods, PAIso-seq1 and PAIso-seq2 to comprehensively analyze the poly(A) tails. PAIso-seq1 is currently the most sensitive poly(A) tail analysis strategy, but limited in catching RNA with very short or no poly(A) tails^1^, while PAIso-seq2 provides complete information about the tail including non-A residues at 5′-end, internal, and 3′-end regardless of their polyadenylation status but limited in sensitivity^36^.

We applied the PAIso-seq1 method to analyze the pig oocytes at germinal vesicle (GV), metaphase I (MI), and metaphase II (MII) stages, as well as preimplantation embryos at the 1-cell (1C), 2-cell (2C), 4-cell (4C), 8-cell (8C), morula (MO) and blastocyst (BL) stages with adequate reproducibility (Extended Data Fig. 1a, b). An overall survey of the data revealed the mRNA poly(A) tail length to be highly dynamic and exhibit a fall-rise trend during development (Fig. 1a). To validate this result, we also performed PAIso-seq2 for GV and MII oocytes, as well as 1C and 2C embryos (Extended Data Fig. 1c, d). The PAIso-seq2 data confirmed the global dynamic changes and the fall-rise trend of global poly(A) tail length distribution (Fig. 1c) as well as poly(A) tail length for individual genes (Fig. 1d) during development.

**Fig. 1.**
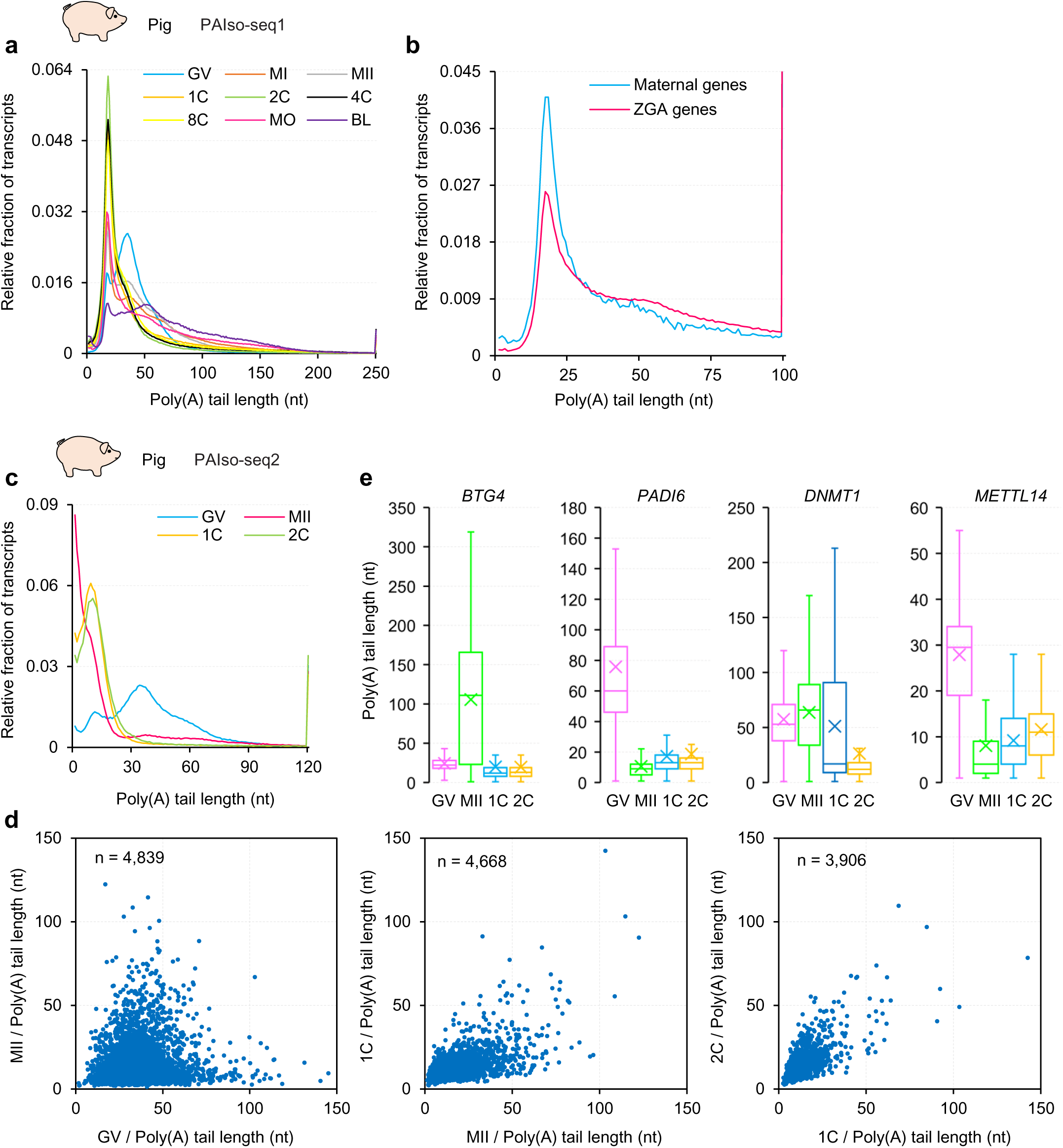
The dynamics of mRNA poly(A) tail length during pig OET. **a.** Histogram of transcriptome-wide poly(A) tails length in pig germinal vesicle (GV), meiosis I (MI), and meiosis II (MII) oocytes, and 1-cell (1C), 2-cell (2C), 4-cell (4C), 8-cell (8C), morula (MO), and blastocyst (BL) embryos measured by PAIso-seq1. Histograms (bin size = 1 nt) are normalized to cover the same area. Transcripts with poly(A) tail of at least 1 nt are included in the analysis. Transcripts with poly(A) tail length greater than 250 nt are included in the 250 nt bin. **b.** Histogram of poly(A) tail length of combined transcripts from maternal genes (n = 2,929) or zygotic genes (n = 2,386) in pig morula embryos measured by PAIso-seq1. Histograms (bin size = 1 nt) are normalized to cover the same area. Transcripts with poly(A) tail of at least 1 nt are included in the analysis. Transcripts with poly(A) tail length greater than 100 nt are included in the 100 nt bin. **c**, Histogram of transcriptome-wide poly(A) tail length in pig GV and MII oocytes, and *in vitro* fertilized 1C and 2C embryos measured by PAIso-seq2. Histograms (bin size = 1 nt) are normalized to cover the same area. Transcripts with poly(A) tail of at least 1 nt are included in the analysis. Transcripts with poly(A) tail length greater than 120 nt are included in the 120 nt bin. **d**, Scatter plot of poly(A) tail length of pig samples at neighboring developmental stages measured by PAIso-seq2. Each dot represents one gene. The poly(A) tail length for each gene is the geometric mean length of all the transcripts with poly(A) tail length of at least 1 nt for the given gene. Genes with at least 10 reads in each sample are included in the analysis. The number of genes included in the analysis is shown on the top left of each graph. **e**, Box plot for the poly(A) tail length of *BTG4*, *PADI6*, *DNMT1,* and *METTL14* in different stage pig samples measured by PAIso-seq2. The “×” indicates mean value. The horizontal bars show the median value. The top and bottom of the box represent the value of 25^th^ and 75^th^ percentile, respectively. Transcripts with poly(A) tail of at least 1 nt for the given gene are included in the analysis.

The global poly(A) tail length dynamic in pigs revealed by PAIso-seq1 and PAIso-seq2 is similar to the observed dynamics of poly(A) tail length during OET in mice and humans^3, 4^, revealing that poly(A) tails undergo global deadenylation during oocyte maturation and global cytoplasmic polyadenylation after fertilization and is conserved across mice, pigs, and humans. In addition, we found the poly(A) tails of mRNA from ZGA are significantly longer than the maternal mRNA in pigs (Fig. 1b), and the maternal transcriptome undergoes both deadenylation (such as *PADI6* and *METTL14*) and cytoplasmic polyadenylation (such as *BTG4* and *DNMT1*) during pig OET (Fig. 1e), as found in mice and humans^3, 4^.

### Highly dynamic non-A residues in mRNA poly(A) tails during pig OET

As in mice and humans^2, 3, 5^, we found that the mRNA poly(A) tails in pig oocytes and early embryos also contain high level of non-A residues. In general, the proportion of poly(A) tail with non-A residues was highly dynamic and showed a rise-fall trend during pig OET in both PAIso-seq1 and PAIso-seq2 datasets (Fig. 2a and Extended Data Fig. 2a). The global and gene-level proportion of non-A residues increased dramatically from GV to MII, then increased gradually until 8C before plunging sharply at the MO stage reaching a level comparable with the GV oocyte stage (Fig. 2a, d and Extended Data Fig. 2a, c). U residues occur in most of the detected transcripts, and G residues less abundant, while C residues least abundant (Fig. 2a and Extended Data Fig. 2a).

**Fig. 2.**
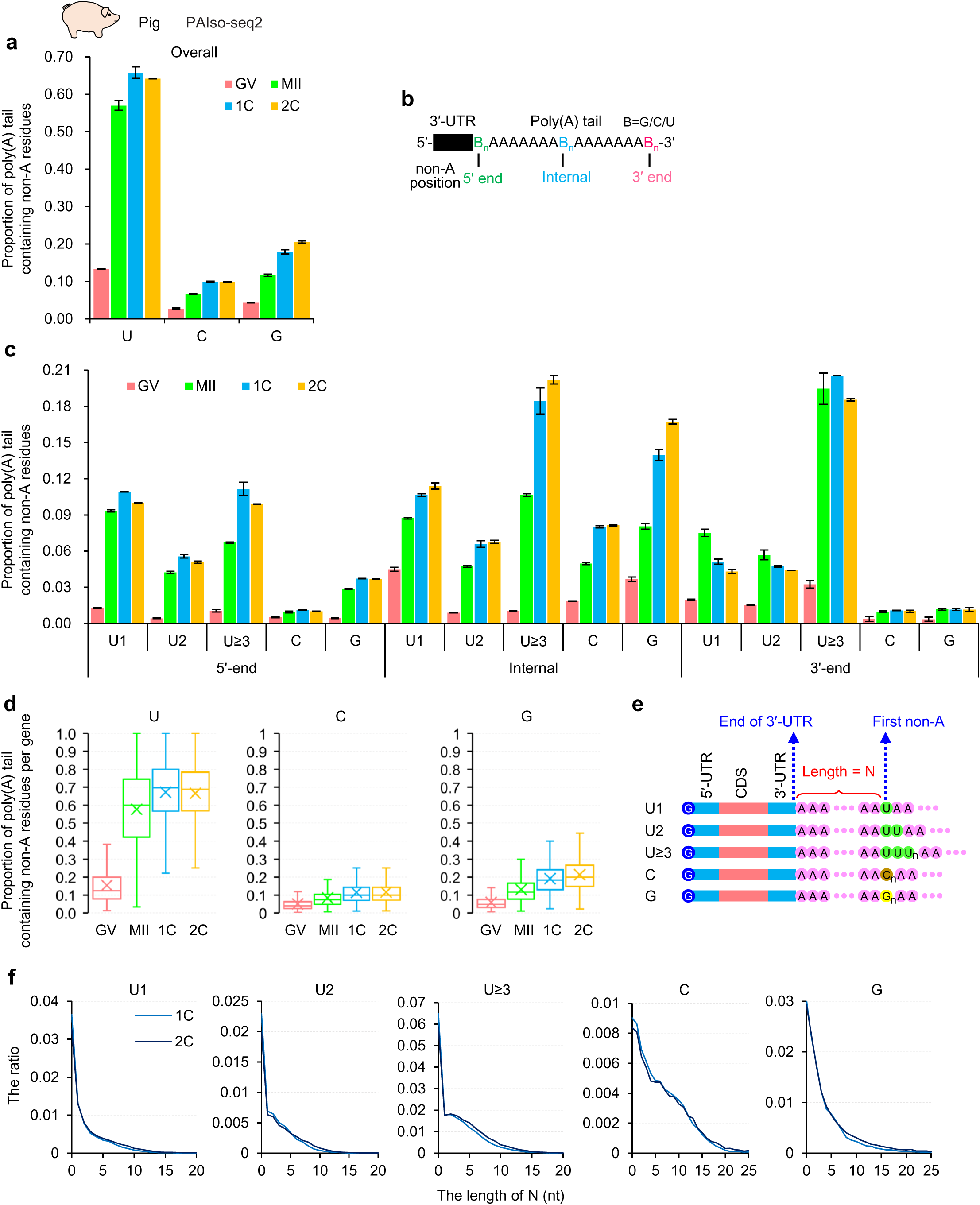
The dynamics of non-A residues during pig OET. **a**, Overall proportion of poly(A) tails containing U, C, and G residues in pig GV and MII oocytes, and 1C and 2C embryos measured by PAIso-seq2. Transcripts with poly(A) tail of at least 1 nt are included in the analysis. Error bars represent the standard error of the mean (SEM) from two replicates. **b**, Diagram depicting mRNA poly(A) tail with 5′-end, internal, and 3′-end non-A residues. **c**, Proportion of poly(A) tails containing U, C, and G residues in the 5′-end, internal, or 3′- end in different stage pig samples measured by PAIso-seq2. The U residues are further divided according to the length of the longest consecutive U (U1, U2, and U≥ 3). Transcripts with poly(A) tail of at least 1 nt are included in the analysis. Error bars represent SEM from two replicates. **d**, Box plot for the proportion of poly(A) tails containing U, C, and G residues of individual genes in different stage pig samples measured by PAIso-seq2. The “×” indicates mean value. The horizontal bars show the median value. The top and bottom of the box represent the value of 25^th^ and 75^th^ percentile, respectively. Genes (n = 3,711) with at least 10 reads and contain poly(A) tails (tail length ≥ 1) in each sample are included in the analysis. **e**, Diagram depicting mRNA poly(A) tail with internal non-A residues. N represents the length of residues between the end of 3′ UTR and the first base of the longest consecutive U, C, or G residues in a poly(A) tail. **f**, Histogram of the length of N and the ratio of U, C, and G residues in pig 1C and 2C embryos measured by PAIso-seq2. The U residues are further divided according to the length of the longest consecutive U (U1, U2, and U≥ 3).

Also as in mice and humans^3, 5^, non-A residues are present not only at the 3′-end but also at the 5′-end and internal regions of the poly(A) tails (Fig. 2b, c and Extended Data Fig. 2b). As U residues often occur consecutively, we further separated the U residues into the three groups (U1, U2, U ≥ 3) based on the maximum length of the consecutive U residues. The three groups of U residues, as well as C and G residues, showed similar dynamics during pig OET at the 5′-end, internal, and 3′-end of poly(A) tails, which is also similar to the overall pattern (Fig. 2a, c and Extended Data Fig. 2a, b), except that U ≥ 3 at the 5′-end and internal, and G in internal increased sharply from MII to 1C (Fig. 2c). The PAIso-seq1 and PAIso-seq2 datasets showed a generally similar pattern, except that PAIso-seq2 can capture the non-A residues at the 3′-end.

We showed that in mice and humans^3, 5^, the non-A residues are a result of cytoplasmic polyadenylation after global deadenylation. The dramatic increases of non-A residues at 5′-end, internal, and 3′-end parts at MI and MII stages (Fig. 2c and Extended Data Fig. 2b), suggest that massive short or fully deadenylated mRNA 3′ tails appear during oocyte maturation^2, 3, 5^, and these tails can be uridylated and re-polyadenylated to produce U residues at the 5′-end and internal positions. Notably, U1 and U2 residues at the 3′-end decreased from MII to 1C stage while they increased slightly in the 5′-end and internal regions, and the level of U3 residues at the 3′-end remained unchanged from MII to 1C stage, while it increased dramatically in the 5′-end and internal parts (Fig. 2c), reflecting that the re-polyadenylation increased greatly after fertilization, as it does in mice and humans^3, 5^. Also as in mice and humans, the C and G residues show a similar pattern regardless of their position (Fig. 2c), implying different mechanisms in controlling the incorporation of G/C residues compared to U residues and different functions of U, C and G residues in poly(A) tails.

Moreover, similar to mice and humans^3, 5^, we discovered that the maternal mRNA in pigs incorporated more non-A residues than ZGA mRNA, especially U residues (Extended Data Fig. 2d), confirming that the dynamics of non-A residues mainly happen in maternal mRNA.

Next, we measured the length, N, between the end of 3′ UTR and the longest consecutive U, C, or G residues in poly(A) tails, for poly(A) tails with U, C, or G residues (Fig. 2e). We found that N is short (with a high level of tails with 0 N value) for all types of poly(A) tails with non-A residues from PAIso-seq1 and PAIso-seq2 datasets (Fig. 2f and Extended Data Fig. 2e), which is similar to mice and humans^3, 5^. This suggests that the non-A residues are added to the short or fully deadenylated tails, consistent with our observation that most of the re-polyadenylation events are on partially degraded mRNA 3′-ends^2^.

In addition, we found that the maternal mRNA with U residues degraded faster than that without U residues from 8C to BL stage (Extended Data Fig. 2f, g), as they do in mice and humans^3, 5^.

Overall, these results revealed the first transcriptome-wide dynamic landscape of mRNA poly(A) tails, including length distribution and non-A residues, during the pig oocyte maturation and preimplantation development.

### Poly(A) tail dynamics during rat OET

Next, we explored mRNA poly(A) tail dynamics during rat OET using PAIso-seq2 with adequate reproducibility (Extended Data Fig. 3). In general, the mRNA poly(A) tail length is also highly dynamic and shows a fall-rise-trend along rat OET, falling sharply during oocyte maturation from GV to MII stage, and then increases gradually after fertilization (Fig. 3a, b). Moreover, as in pigs (Fig. 1b, e), mice, and humans^3, 4^, we found that the poly(A) tails of ZGA mRNA are significantly longer than the maternal mRNA in rats (Fig. 3c), and both deadenylation (such as *Cnot7*, *Ooep*, and *Paid6*) and cytoplasmic polyadenylation (such as *Btg4*) occur to the maternal transcriptome during rat OET (Fig. 3d).

**Fig. 3.**
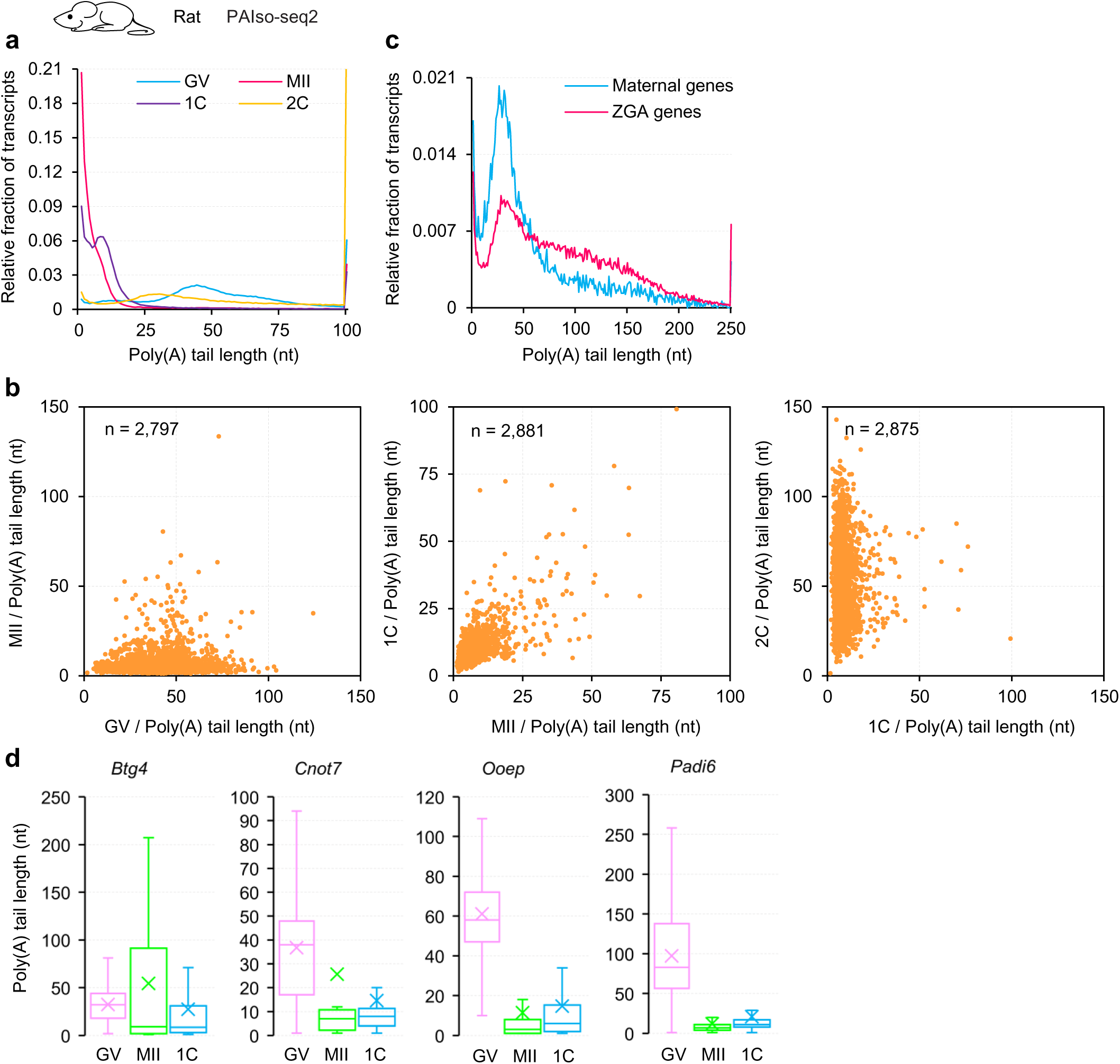
The dynamics of mRNA poly(A) tail length during rat OET. **a**, Histogram of transcriptome-wide poly(A) tail length in rat GV and MII oocytes, and 1C and 2C embryos measured by PAIso-seq2. Histograms (bin size = 1 nt) are normalized to cover the same area. Transcripts with poly(A) tail of at least 1 nt are included in the analysis. Transcripts with poly(A) tail length greater than 100 nt are included in the 100 nt bin. **b**, Scatter plot of poly(A) tail length of rat samples at neighboring developmental stages measured by PAIso-seq2. Each dot represents one gene. The poly(A) tail length for each gene is the geometric mean length of all the transcripts with a poly(A) tail length of at least 1 nt for the given gene. Genes with at least 10 reads in each sample are included in the analysis. The number of genes included in the analysis is shown on the top left of each graph. **c.** Histogram of poly(A) tails length of combined transcripts from maternal genes (n = 1,519) or zygotic genes (n = 1,699) in rat 2C embryos measured by PAIso-seq2. Histograms (bin size = 1 nt) are normalized to cover the same area. Transcripts with poly(A) tail of at least 1 nt are included in the analysis. Transcripts with poly(A) tail length greater than 250 nt are included in the 250 nt bin. **d**, Box plot for the poly(A) tail length of *Btg4*, *Cnot7*, *Ooep,* and *Paid6* in different stage rat samples measured by PAIso-seq2. The “×” indicates mean value. The horizontal bars show the median value. The top and bottom of the box represent the value of 25^th^ and 75^th^ percentile, respectively. Transcripts with poly(A) tail of at least 1 nt for the given gene are included in the analysis.

As in mice, pigs, and humans, the mRNA poly(A) tails during rat OET also contain non-A residues. In general, the proportion of poly(A) tails with non-A residues were highly dynamic during rat OET, increasing dramatically until 1C stage, then decreasing sharply at 2C stage, not only for the transcriptome (Fig. 4a) but also for individual genes (Fig. 4c). Among the U, C, and G residues, U residues are the most abundant, occurring in up to 50% of detected transcripts, followed by the G residues, with C residues being the least abundant (Fig. 4a).

**Fig. 4.**
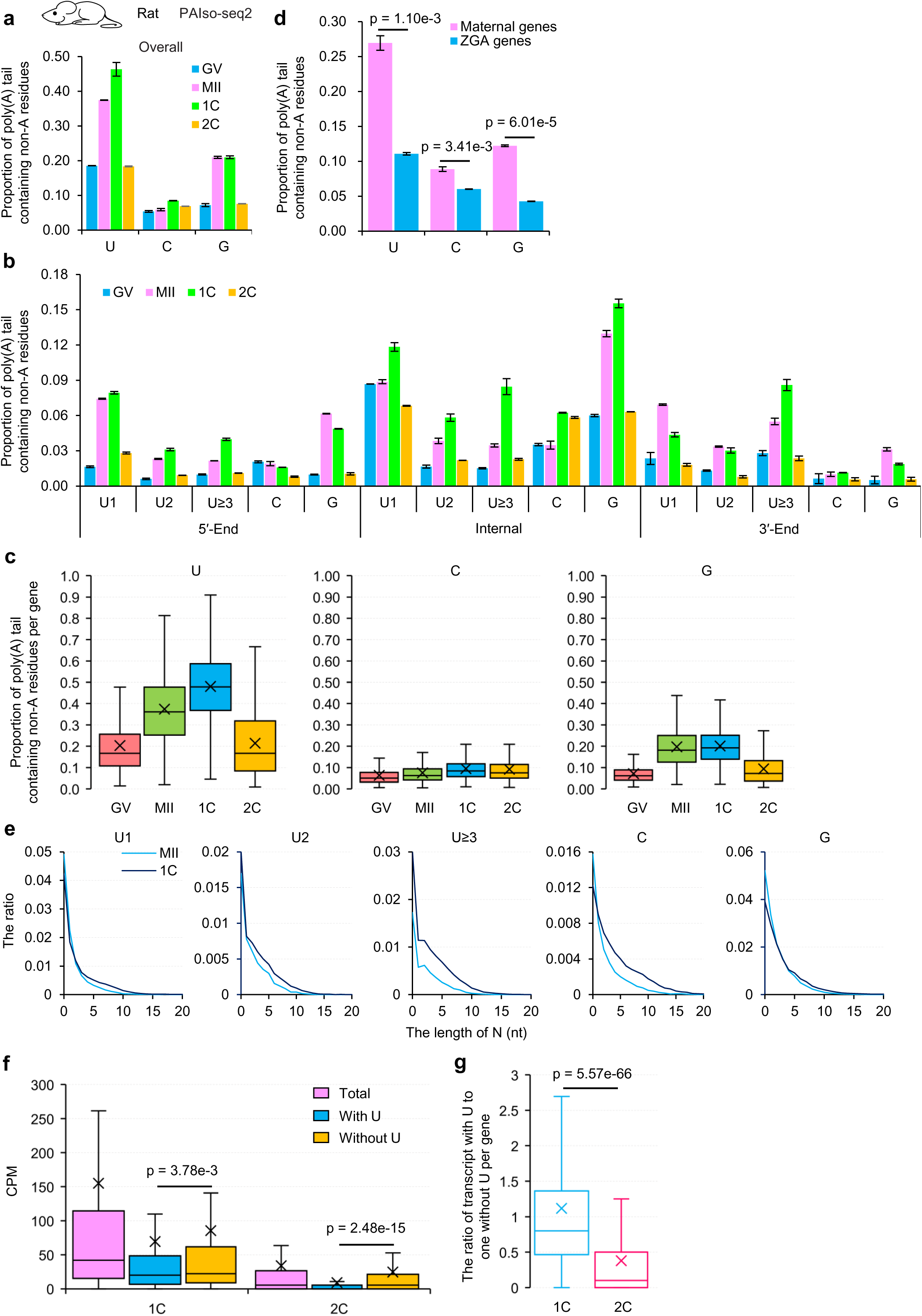
The dynamics of non-A residues during rat OET. **a**, Overall proportion of poly(A) tails containing U, C, and G residues in rat GV and MII oocytes, and 1C and 2C embryos measured by PAIso-seq2. Transcripts with poly(A) tail of at least 1 nt are included in the analysis. Error bars represent SEM from two replicates. **b**, Proportion of poly(A) tails containing U, C, and G residues in the 5′-end, internal, or 3′- end in different stages of rat samples measured by PAIso-seq2. The U residues are further divided according to the length of the longest consecutive U (U1, U2, and U≥ 3). Transcripts with poly(A) tail of at least 1 nt are included in the analysis. Error bars represent SEM from two replicates. **c**, Box plot for the proportion of poly(A) tails containing U, C, and G residues of individual genes in different stages of rat samples measured by PAIso-seq2. Genes (n = 1,736) with at least 10 reads and contain poly(A) tails (tail length ≥ 1 nt) in each sample are included in the analysis. **d**, Overall proportion of transcripts containing U, C, or G residues for combined transcripts from maternal genes (n = 1,519) or zygotic genes (n = 1,699) in rat 2C embryos measured by PAIso-seq2. Transcripts with poly(A) tail of at least 1 nt are included in the analysis. Error bars represent SEM from two replicates. *P-*value is calculated by Student’s *t-*test. **e**, Histogram of the length of N and the ratio of U, C, and G residues in rat MII oocytes and 1C embryos measured by PAIso-seq2. The U residues are further divided according to the length of the longest consecutive U (U1, U2, and U≥ 3). **f**, Box plot for the expression level of total transcripts, transcripts with U residues, and transcripts without U residues for each maternal gene in rat 1C and 2C embryos measured by PAIso-seq2. *P*-value is calculated by Student’s *t*-test. **g**, Box plot for the ratio of number of transcripts with U residues to those without U residues for each maternal gene in rat 1C and 2C embryos measured by PAIso-seq2. *P-*value is calculated by Student’s *t*-test. For all box plots, the “×” indicates mean value. The horizontal bars show the median value. The top and bottom of the box represent the value of 25^th^ and 75^th^ percentile, respectively.

Similarly, non-A residues were also present at the 3′-end, 5′-end, and interior of the poly(A) tails (Fig. 4b). As U residues were also found consecutively, we further separated the U residues into three groups (U1, U2, U≥3) based on the maximum length of the consecutive U residues. The three groups of U residues, as well as C and G residues, showed similar dynamics patterns during rat OET in 5′-end, internal, and 3′-end of poly(A) tails, which is also similar to the overall pattern (Fig. 4a, b).

Similar to the patterns in mice, pigs, and humans^3, 5^, we also found dramatic increases of non-A residues at 5′-end, internal, and 3′-end segments from the GV to MII stages (Fig. 4b), suggesting that massive short or fully deadenylated mRNA 3′ tails appeared during oocyte maturation^2, 3, 5^ and then uridylated and re-polyadenylated to produce U residues at the 5′-end and internal positions. Notably, the abundance of U1 residues at the 3′-end decreased from the MII to 1C stage while they increased in the 5′-end and internal parts, and the pattern of U2 residues at the 3′-end remain unchanged from the MII to 1C stage, while they increased in the 5′-end and internal parts (Fig. 4b), reflecting that the re-polyadenylation increases greatly after fertilization, which is similar to mouse, human^3, 5^, and pig. However, the pattern of U3 residues is similar in 5′-end, internal, and 3′-end parts. Moreover, the C and G residues in poly(A) tails showed a similar pattern regardless of its position at the 5′-end, the interior, or the 3′-end (Fig. 4b), implying different mechanisms for controlling the incorporation of G/C residues compared to U residues, and different functions of U, C, and G residues in poly(A) tails.

Moreover, we also discovered maternal mRNA incorporated more non-A residues than ZGA mRNA, especially U residues (Fig. 4d), confirming that the dynamics of non-A residues mainly happen in maternal mRNA.

In addition, we also found that the N is short (with a high level of tails with 0 N value) for all types of poly(A) tails with non-A residues (Fig. 4e), suggesting that the non-A residues are added to the short deadenylated tails or fully deadenylated tails, consistent with our observation that most of the re-polyadenylation events are on partially degraded mRNA 3′-ends^2^. In addition, maternal mRNA with U residues degraded faster than those without U residues (Fig. 4f, g). These results revealed the first transcriptome-wide dynamic landscape of mRNA poly(A) tails during the rat OET.

Overall, we revealed some conserved characteristics of poly(A) tail dynamic during OET across all four species: a rise-fall pattern in poly(A) tail length distribution; a fall-rise pattern in the ratio of poly(A) tail with non-A residues; abundant consecutive U residues in poly(A) tails of maternal mRNA; and increasing re-polyadenylation activity after fertilization.

### Abnormal ZGA activation and maternal RNA deadenylation in mouse and rat parthenogenetic embryos

To further explore the poly(A) tail dynamics during mammalian OET, we paid attention to parthenogenetic embryos. We performed parthenogenetic activation with mouse and rat oocytes followed by PAIso-seq2 analysis at 2C stage (Pa-2C) (Fig. 5a and Extended Data Fig. 4). As a result, we found an insufficient activation of ZGA and impaired maternal mRNA decay in Pa-2C embryos, as the expression levels of ZGA genes were generally lower and the levels of maternal genes were globally higher in Pa-2C than fertilized 2-cell (2C) in both mice and rats (Fig. 5b, c, g, h). Considering that maternal mRNA decay is mainly deadenylation-dependent, we wondered whether the poly(A) tail length and non-A residues of maternal mRNA were abnormal in Pa-2C. Interestingly, we found the poly(A) tail length distribution for maternal genes showed a right shift of the distribution pattern in Pa-2C compared to 2C in mice and rats (Fig. 5d, i), reflecting that the maternal mRNA deadenylation in Pa-2C was impaired. Moreover, the ratio of poly(A) tails with non-A residues of maternal genes was higher in Pa-2C not only at the transcript but also at the gene level in mice and rats (Fig. 5e, f, j, k), indicating the maternal mRNA deadenylation was impaired because the maternal mRNA poly(A) tail incorporated more non-A residues than ZGA mRNA in mice and humans^3, 5^.

**Fig. 5.**
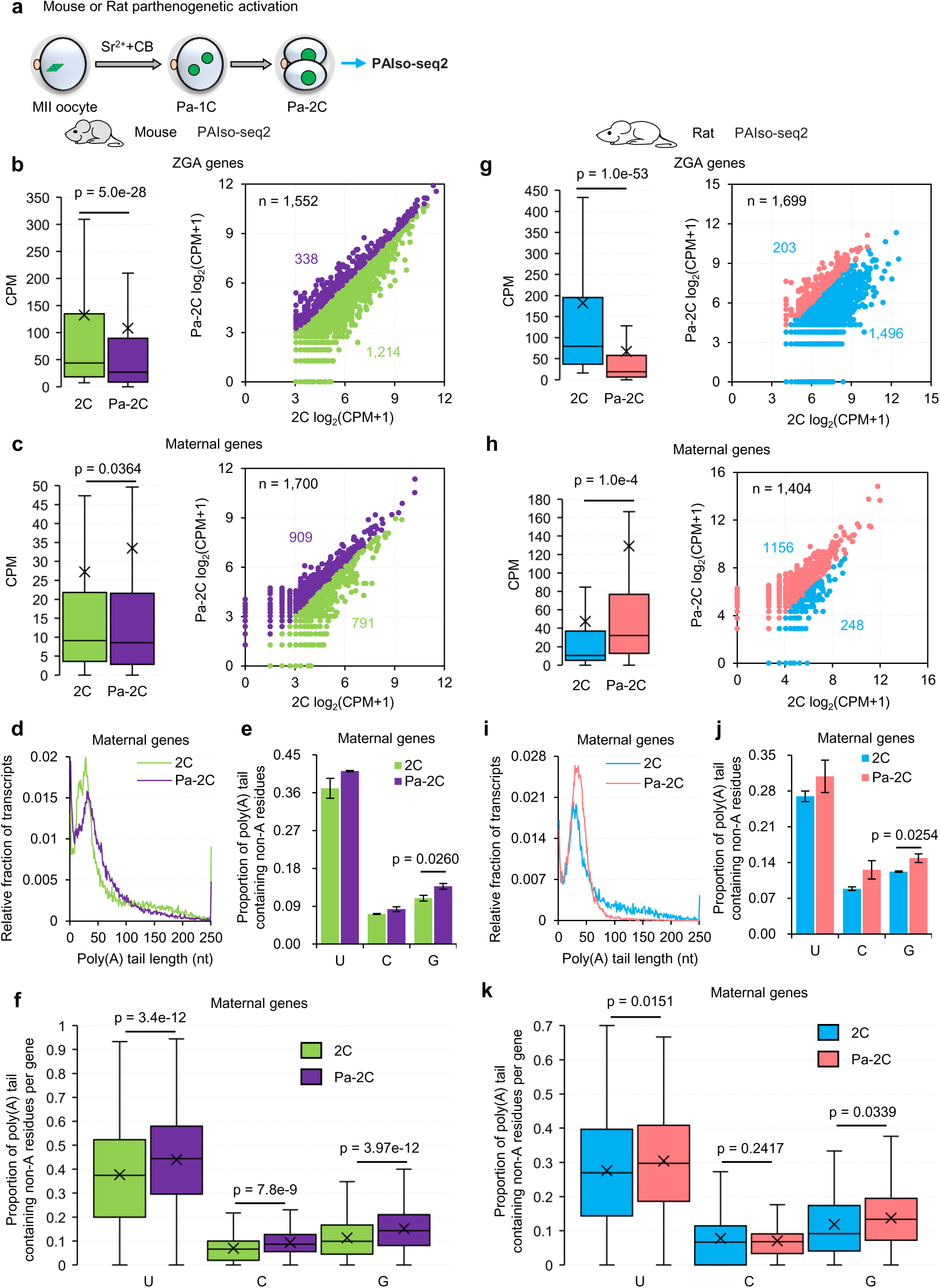
Abnormal ZGA activation and maternal RNA deadenylation in mouse and rat parthenogenetic 2C embryos. **a**, Flowchart of collection of parthenogenetic 2C (Pa-2C) embryos in mouse and rat. **b, c, g, h**, Box plot (left) or scatter plot (right) for the transcriptional level of ZGA genes (**b**, **g**) or maternal genes (**c**, **h**) in mouse (**b**, **c**) or rat (**g**, **h**) fertilized 2C (2C) and Pa-2C embryos measured by PAIso-seq2. For scatter plot, each dot represents one gene; Dots in purple (**b**, **c**) or in red (**g**, **h**) are genes with a higher expression level in Pa-2C, while dots in green (**b**, **c**) or in blue (**g**, **h**) are genes with a higher expression level in Pa-2C; and the number of genes included in the analysis is shown on the top left of each graph. Gene numbers for the box plot are the same as the associated scatter plot with the same color. *P*-value is calculated by Student’s *t-*test. **d, i**, Histogram of poly(A) tail length of maternal mRNA in mouse (**d**, n = 1,700) or rat (**i**, n = 1,404) 2C and Pa-2C embryos measured by PAIso-seq2. Histograms (bin size = 1 nt) are normalized to cover the same area. Transcripts with poly(A) tail of at least 1 nt are included in the analysis. Transcripts with poly(A) tail length greater than 250 nt are included in the 250 nt bin. **e, j**, Proportion of poly(A) tails containing U, C, and G residues in maternal genes in mouse (**e**, n = 1,700) or rat (**j**, n = 1,404) 2C and Pa-2C embryos measured by PAIso-seq2. Transcripts with poly(A) tail of at least 1 nt are included in the analysis. Error bars represent SEM from two replicates. *P*-value is calculated by Student’s *t*-test. **f, k**, Box plot for the proportion of poly(A) tails containing U, C, and G residues of individual maternal genes in mouse (**f**) or rat (**k**) 2C and Pa-2C embryos measured by PAIso-seq2. Genes (**f**, n = 354; **k**, n = 178) with at least 10 reads and contain poly(A) tails in each sample are included in the analysis. *P*-value is calculated by Student’s *t-*test. For all box plots, the “×” indicates mean value. The horizontal bars show the median value. The top and bottom of the box represent the value of 25^th^ and 75^th^ percentile, respectively.

### Inhibition of ZGA leads to blocked deadenylation of maternal mRNA in human and mouse embryos

ZGA inhibition via α-amanitin treatment is reported to block the maternal mRNA decay^37, 38^, however, the underlying mechanisms are still unknown. We thus wondered if this blockage is due to defective maternal mRNA deadenylation. We took advantage of PAIso-seq1 to detect mRNA poly(A) tail length and non-A residues in mouse and human embryos with α-amanitin treatment (Fig. 6a, g and Extended Data Fig. 5). As expected, the ZGA was inhibited upon α-amanitin treatment in both mouse and human embryos, as the expression level of ZGA genes decreased dramatically (Fig. 6b, h). In addition, we found the level of maternal genes was globally higher upon α-amanitin treatment both in mouse and human embryos (Fig. 6c, i), indicating that ZGA inhibition leads to impaired maternal mRNA decay. Interestingly, we found the maternal mRNA poly(A) tail length distribution pattern shifted to the right upon α-amanitin treatment in mouse and human embryos (Fig. 6d, j), indicating that the maternal mRNA deadenylation upon ZGA inhibition was impaired. Moreover, we discovered that the proportion of poly(A) tails with non-A residues increased not only at transcript but also at the individual gene level of maternal genes upon α-amanitin treatment in mouse and human embryos (Fig. 6e, f, k, l), further confirming an impairment of maternal mRNA deadenylation because maternal mRNA incorporated more non-A residues in mice and humans^3, 5^. These results indicated that ZGA inhibition leads to blocked deadenylation of maternal mRNA.

**Fig. 6.**
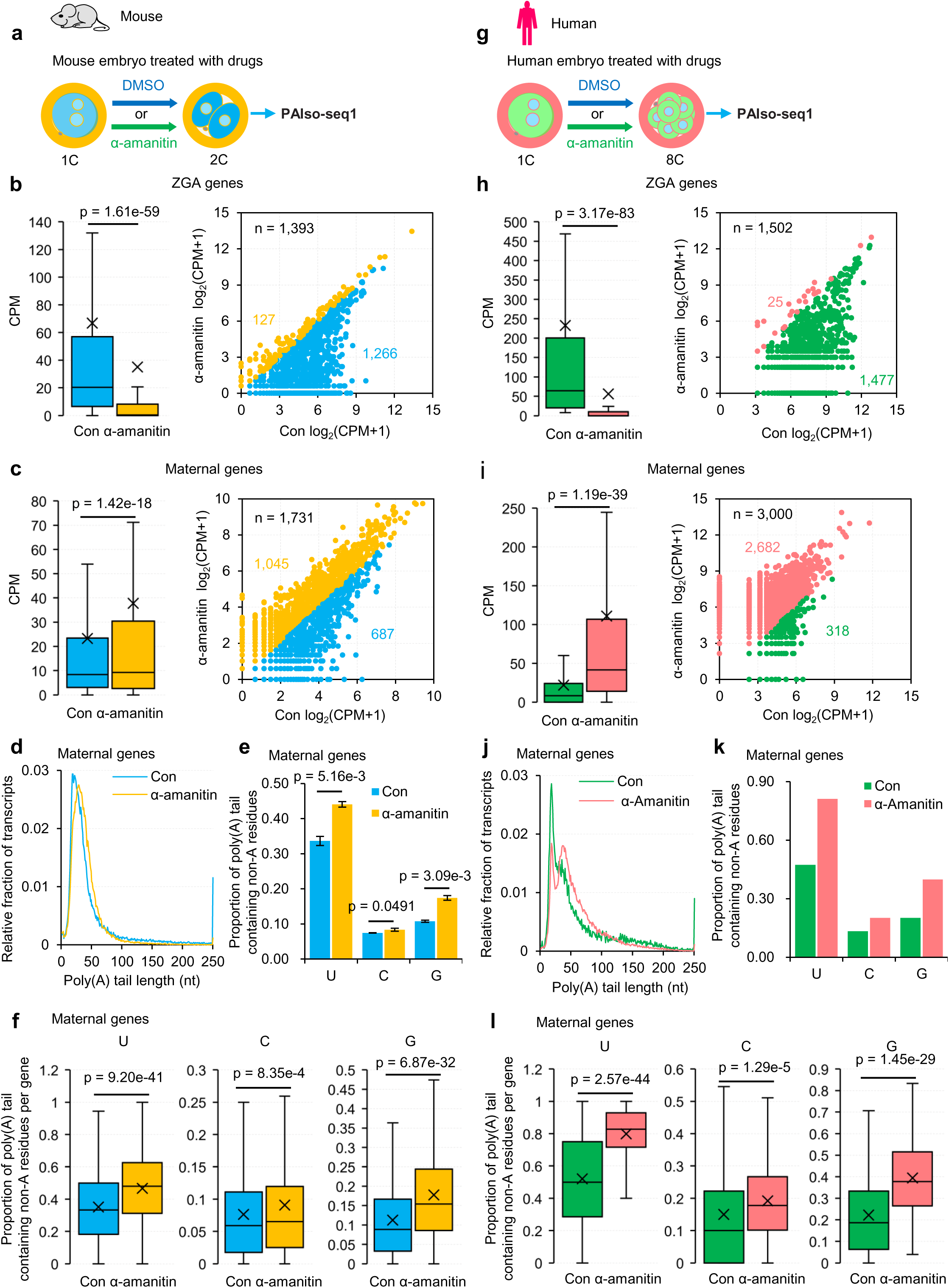
Inhibition of ZGA in human or mouse embryos results in impaired maternal mRNA deadenylation. **a, g**, Flowchart of collection of embryos with ZGA inhibition by α-amanitin treatment in mouse (**a**) and human (**g**). **b, c, h, i**, Box plot (left) or scatter plot (right) for the transcriptional level of ZGA genes (**b**, **h**) or maternal genes (**c**, **i**) in mouse 2C (**b**, **c**) or human (**h**, **i**) 8C embryos with or without α-amanitin treatment measured by PAIso-seq1. For scatter plot, each dot represents one gene; Dots in orange (**b**, **c**) or in red (**h**, **i**) are genes with a higher expression level in α-amanitin treated embryos, while dots in blue (**b**, **c**) or in green (**h**, **i**) are genes with higher expression level in control embryos; and the number of genes included in the analysis is shown on the top left of each graph. Gene numbers for the box plots are the same as for the associated scatter plot with the same color. *P*-value is calculated by Student’s *t*-test. **d, j**, Histogram of poly(A) tail length of maternal mRNA in mouse 2C (**d**, n = 1,731) or human (**j**, n = 3,000) 8C embryos with or without α-amanitin treatment measured by PAIso-seq1. Histograms (bin size = 1 nt) are normalized to cover the same area. Transcripts with poly(A) tail of at least 1 nt are included in the analysis. Transcripts with poly(A) tail length greater than 250 nt are included in the 250 nt bin. **e, k**, Proportion of poly(A) tails containing U, C, and G residues in maternal genes in mouse 2C (**e**, n = 1,731) or human (**k**, n = 3,000) 8C embryos with or without α-amanitin treatment measured by PAIso-seq1. Transcripts with poly(A) tail of at least 1 nt are included in the analysis. Error bars represent SEM from two replicates. *P*-value is calculated by Student’s *t*-test. **f, l**, Box plot for the proportion of poly(A) tails containing U, C, and G residues of individual maternal genes in mouse 2C (**f**) or human (**l**) 8C embryos with or without α-amanitin treatment measured by PAIso-seq1. Genes (**f**, n = 809; **k**, n = 329) with at least 10 reads and contain poly(A) tails (tail length ≥ 1 nt) in each sample are included in the analysis. *P*-value is calculated by Student’s *t*-test. For all box plots, the “×” indicates mean value. The horizontal bars show the median value. The top and bottom of the box represent the value of 25^th^ and 75^th^ percentile, respectively.

### Differences in the duration time of U residues and the length of continuous U residues across four mammals

After identifying several conserved dynamics of poly(A) tail length and non-A residues during OET across species, we wondered about possible differences. Considering U residues are the most enriched non-A residues in maternal mRNA poly(A) tails, we compared the dynamics of U residues during oocyte maturation and early embryonic development across the four species from PAIso-seq1 (Fig. 7a) and PAIso-seq2 (Fig. 7b) datasets. The results showed that the rise-fall trend of the ratio of poly(A) tails with U residues is conserved across the four mammals, yet the duration time of U residues diverges (Fig. 7a, b). In detail, the proportion of poly(A) tails with U residues drop to a level as low as the GV stage at 2C in rats, at 4C in mice, at 8C in humans, and at morula in pigs (Fig. 7a, b).

**Fig. 7.**
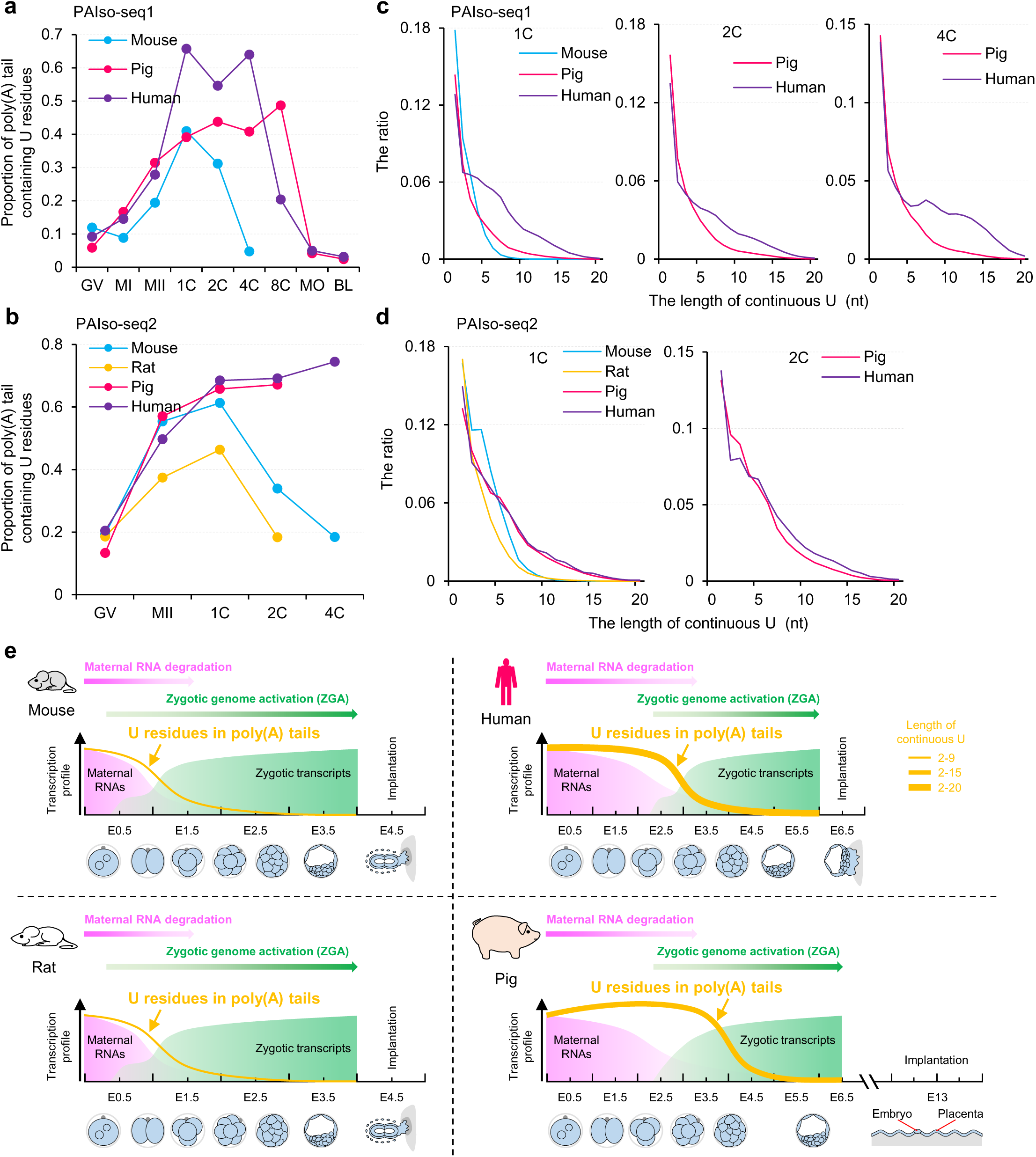
Conservation and divergence of the length of consecutive U residues and the duration time of U residues in mice, rats, pigs and humans. **a, b**, Proportion of poly(A) tails containing U residues during OET in mouse, rat, pig, and human f measured by PAIso-seq1 (**a**) or PAIso-seq2 (**b**). Transcripts with poly(A) tail of at least 1 nt are included in the analysis. **c, d**, Histogram of the length of continuous U and the ratio of U residues in 1C, 2C, and 4C embryos in mouse, rat, pig, and human measured by PAIso-seq1 (**c**) or PAIso-seq2 (**d**). Transcripts with poly(A) tail of at least 1 nt are included in the analysis. **e**, Model showing maternal RNA decay, waves of ZGA, and dynamic of poly(A) tail U residues during preimplantation development in mouse, rat, pig, and human.

Another interesting aspect of U residues is the consecutive U residues, then we analyzed the length of consecutive U residues across the four mammals. We found the length can be up to 10 nt in mouse and rat embryos, and up to 20 nt in pig and human embryos (Fig. 7c, d). In addition, the length of consecutive U residues is generally longer in humans than in pigs (Fig. 7c).

These results revealed two differences in poly(A) tail dynamics during OET across mice, rats, pigs, and humans that the duration time of U residues and the length of consecutive U residues are different among the four mammals (Fig. 7e).

## Discussion

The OET in mammals is very important for full-term development, and the maternal mRNA plays a dominant role because transcription is silent before ZGA. Here, we revealed both conservation and divergence in poly(A) tail dynamics during OET across mice, rats, pigs, and humans. The dynamic change of poly(A) tail length distribution and the proportion of poly(A) tails with non-A residues are conserved; a higher level of non-A residues in poly(A) tails of maternal mRNA than in zygotic genome activation (ZGA) mRNA is conserved; and impedance of maternal mRNA deadenylation by parthenogenetic activation and ZGA inhibition is conserved. In contrast, the length of consecutive U residues and the duration time of U residues in poly(A) tail diverges across the four mammals. Our finding raises new clues for the understanding of OET across different mammal species.

The decay of maternal mRNA has been shown to be necessary for successful ZGA^31, 32^. Here we found impaired ZGA would inhibit maternal mRNA deadenylation (Fig. 6). This result showed a positive feedback relationship between maternal mRNA decay and ZGA, which are the two core molecular events of OET. In zebrafish, the minor ZGA produces miR-430 for degradation of a large pool of maternal RNA through accelerated deadenylation^39^. Nevertheless, the mechanisms underlying this positive feedback loop in mammals warrants further investigation, including how ZGA and maternal mRNA deadenylation regulate each other, whether some transcripts from ZGA promote maternal mRNA deadenylation, and the degree of conservation of mechanisms or genes involved across different mammals is unknown.

The lengths of consecutive U residues are longer in pigs and humans than those in mice and rats (Fig. 7e). This phenomenon is very interesting. The function and regulation of this divergence awaits further investigation. It is worth studying whether this is an adaptive evolutionary mechanism to accommodate prolonged early embryonic development in humans and pigs than the rodents.

We found that maternal mRNA deadenylation is impaired in parthenogenetic embryos, reflecting roles of paternal contribution in maternal mRNA deadenylation after fertilization, suggesting interesting interaction between the maternal and paternal contribution to the zygotes. The mechanism of paternal contribution to maternal mRNA deadenylation warrants future exploration. Parthenogenetic embryos are not the only embryos without paternal contribution. The cloned embryos derived from somatic cell nuclear transfer are also no paternal contribution. The cloned embryos are well-known to be of impaired ZGA as well as of lower developmental potential. Therefore, it is very likely that the maternal RNA deadenylation in cloned embryos is also impaired due to the lack of paternal contribution, which represents a post-transcriptional regulation barrier for cloned embryos. Therefore, it will be interesting to investigate this potential cloning barrier in the future and it points a new direction to improve cloning efficiency by overcoming this potential barrier.

## Material and Methods

### Animal care and use

Mice were purchased from Beijing Vital River Laboratory Animal Technology Co., Ltd., and used in compliance with the guidelines of the Animal Care and Use Committee of the Institute of Genetics and Development Biology, Chinese Academy of Sciences. Rats were purchased from Beijing Vital River Laboratory Animal Technology Co., Ltd., and used in compliance with the guidelines of the Animal Care and Use Committee of the Institute of Zoology, Chinese Academy of Sciences. All pig experiments followed the guidelines of Animal Care and Use Committee of the Northeast Agricultural University.

### Porcine oocytes and embryos collection

Ovaries were collected from a slaughterhouse and transported to the lab in 0.9% saline at 37°C. Follicles 3 to 5 millimeters in diameter were aspirated to obtain cumulus-oocyte complexes, which were cultured 42 hours for maturation in an atmosphere of 5% CO_2_ and 95% air at 39°C. MI oocytes and MII oocytes with a clearly extruded first polar body were collected at 36 and 42 hours, respectively.

For IVF, the spermatozoa and cumulus-free oocytes were washed three times in a modified Tris-buffered medium (mTBM). Approximately 30 oocytes were inseminated in 50 μL drops of mTBM at a final sperm concentration of 3 × 10^5^ /mL for 6 hours. Then embryos were washed and cultured in PZM-3 medium for 7 days in an atmosphere of 5% CO_2_ and 95% air at 39°C. IVF embryos at 1C, 2C, 4C, 8C, Mo, and blastocyst (BL) stages were collected 6, 24, 48, 72, 108, and 168 hours after fertilization respectively.

### Rat oocytes and embryos collection

Female rats were superovulated by intraperitoneal injection of 150 IU/kg of pregnant mare serum gonadotropin (PMSG, ProSpec, HOR-272) and 300 IU/kg of human chorionic gonadotropin (hCG, ProSpec, HOR-250) 51 to 52 hours later. GV oocytes were collected 48 hours post-PMSG injection without hCG administration. MII oocytes were collected 14 hours post hCG injection. Fertilized 1C and 2C embryos were collected in M2 medium (Sigma, M7167) containing MG-132 (Sigma, 474790) 20 and 60 hours post hCG injection and mating with male rats, respectively.

Parthenogenetic 2C (Pa-2C) embryos were acquired as follows: MII oocytes were placed in M2 medium with MG-132, then activation was induced by transfer of oocytes to R1EMC containing 150 µM Butyrolactone I (Affiniti R.P.L, Mamhead, UK) and 5 µM Cytochalasin B (Abcam) for 2 hours. Activated oocytes were transferred to R1EMC containing 5 µM Cytochalasin B for 4 hours and finally placed in the M2 medium with MG-132 for another 48 hours before Pa-2C collection.

### Mouse oocytes and embryos collection

Eight-week-old female mice were super-ovulated by consecutive injection of pregnant mare serum gonadotropin (PMSG, ProSpec, HOR-272) and human chorionic gonadotropin (hCG, ProSpec, HOR-250). Oocytes were collected from the oviduct 13 to15 hours post hCG injection and cultured in M16 medium (Sigma, MR-016).

For *in vitro* fertilization, sperm was obtained from the cauda epididymis and transferred to capacitate for 1 hour in human tubal fluid (HTF) medium (Sage, Bedminster, NJ, USA) in an atmosphere of 5% CO_2_ and 95% air at 37°C. Oocytes and sperm were co-incubated in HTF medium for another 4 hours. Fertilized oocytes were washed several times in M16 and transferred to 50 μL drops of M16 medium covered with paraffin oil. The embryos were further cultured in M16 in an atmosphere of 5% CO_2_ and 95% air at 37°C. After another 24 hours, *in vitro* fertilized 2-cell embryos were collected.

For parthenogenetic activation, MII oocytes were placed in M2 medium, then activation was induced by transfer of oocytes to calcium-free CZB containing 5 mg/mL cytochalasin B and 10 mM SrCl_2_ for 5 to 6 hours. Next, activated oocytes were transferred to M16 at 37°C with 5% CO_2_. After another 24 hours, parthenogenetic 2-cell embryos were collected. For drug treatment, the α-amanitin (Sigma, 94072) was dissolved in M16 medium. 1-cell embryos were cultured in a medium containing α-amanitin or DMSO as a control. The control and α-amanitin-treated embryos were counted or collected at the 2-cell stage.

### PAIso-seq1 / PAIso-seq2 library construction and data processing

Libraries for PAIso-seq1 were constructed as described previously^1^. PAIso-seq2 libraries were constructed following the PAIso-seq2 protocol described in another study^36^. The PAIso-seq1 and PAIso-seq2 data generated in this study were processed following the methods described^3–5^.

### Analysis of maternal and zygotic genes

Read counts of each gene were summarized using *featureCounts*. The maternal and zygotic genes were defined using the PAIso-seq1 or PAIso-seq2 data following the published strategy with minor modification^40^. In brief, *edgeR* was used for differential expression analysis^41^. The maternal genes were defined by protein-coding genes with showing 4-fold enrichment in GV oocytes compared to 2C (mouse and rat), 8C (human), or marula (pig) embryos at the transcript level (*p* < 0.05), while the zygotic genes were defined by protein-coding genes with showing 4-fold enrichment in 2C (mouse and rat), 8C (human) or marula (pig) embryos compared to GV oocytes at the transcript level (*p* < 0.05).

### Statistical analyses

Statistical analyses [mean ± standard error of the mean (SEM)] were performed in Excel. Levels of significance were calculated with Student’s *t*-tests.

### Data availability

The ccs data in bam format from PAIso-seq1 and 2 experiments will be available at Genome Sequence Archive hosted by National Genomic Data Center. Custom scripts used for data analysis will be available upon request.

## Acknowledgements

We thank Shuang Wu for her technical assistance in mouse superovulation and collection of oocytes and embryos. We thank Hu Nie for his technical assistance in bioinformatic analysis. This work was supported by the National Key Research and Development program of China (2016YFA0100200, 2018YFA0107001), the Strategic Priority Research Program of the Chinese Academy of Sciences (XDA24020203), National Natural Science Foundation of China (31970588, 32170606), Natural Science Foundation of Heilongjiang province (YQ2020C003), the China Postdoctoral Science Foundation (2020M670516, 2020T130687), the State Key Laboratory of Molecular Developmental Biology, Heilongjiang Touyan Innovation Team Program and the Fundamental Research Funds of Shandong University.

## Author Contributions

Yusheng Liu, Falong Lu and Jiaqiang Wang conceived the project and designed the study. Yusheng Liu constructed the library of the PAIso-seq1 and PAIso-seq2. Yusheng Liu, Yiwei Zhang, Falong Lu and Jiaqiang Wang analyzed the sequencing data. Junxue Jin and Zhonghua Liu collected pig oocytes and embryos. Leyun Wang and Wei Li collected rat oocytes and embryos. Yusheng Liu collected and performed drug treatment on mouse embryos. Chuanxin Zhang and Zhenzhen Hou collected and performed drug treatment on human embryos. Yusheng Liu and Jiaqiang Wang organized all figures. Yusheng Liu, Falong Lu and Jiaqiang Wang supervised the project. Yusheng Liu, Falong Lu and Jiaqiang Wang wrote the manuscript with the input from the other authors.

---

**Extended Data Fig. 1.**
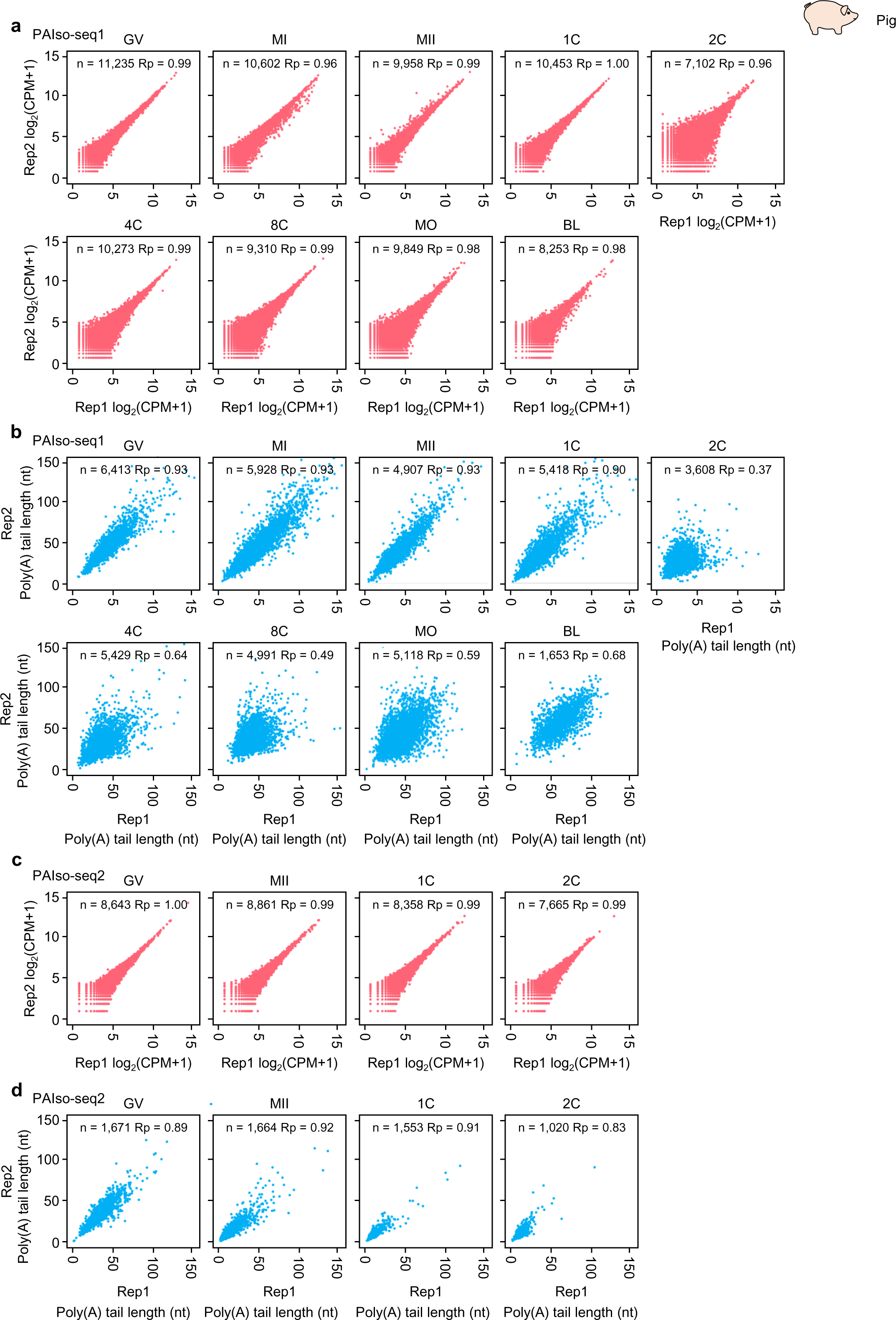
The reproducibility of pig samples. **a**, Scatter plot showing the Pearson correlation of gene expression between two replicates for pig germinal vesicle (GV), metaphase I (MI), and metaphase II (MII) oocytes, and 1-cell (1C), 2-cell (2C), 4-cell (4C), 8-cell (8C), morula (MO), and blastocyst (BL) embryos measured by PAIso-seq1. Each dot represents one gene. Pearson’s correlation coefficient (Rp) and the number of genes included in the analysis are shown on the top of each graph. **b**, Scatter plot showing the Pearson correlation of poly(A) tail length between two replicates for pig GV, MI, and MII oocytes, and 1C, 2C, 4C, 8C, MO, and BL embryos measured by PAIso-seq1. Each dot represents one gene. The poly(A) tail length for each gene is the geometric mean length of all the transcripts with poly(A) tail length of at least 1 nt for the given gene. Genes with at least 20 reads in each sample are included in the analysis. Pearson’s correlation coefficient (Rp) and the number of genes included in the analysis are shown on the top of each graph. **c**, Scatter plot showing the Pearson correlation of gene expression between two replicates for pig GV, MI, and MII oocytes, and 1C and 2C embryos measured by PAIso-seq2. Each dot represents one gene. Pearson’s correlation coefficient (Rp) and the number of genes included in the analysis are shown on the top of each graph. **d**, Scatter plot showing the Pearson correlation of poly(A) tail length between two replicates for pig GV, MI, and MII oocytes, and 1C and 2C embryos measured by PAIso-seq2. Each dot represents one gene. The poly(A) tail length for each gene is the geometric mean length of all the transcripts with a poly(A) tail of at least 1 nt for the given gene. Genes with at least 20 reads in each sample are included in the analysis. Pearson’s correlation coefficient (Rp) and the number of genes included in the analysis are shown on the top of each graph.

**Extended Data Fig. 2.**
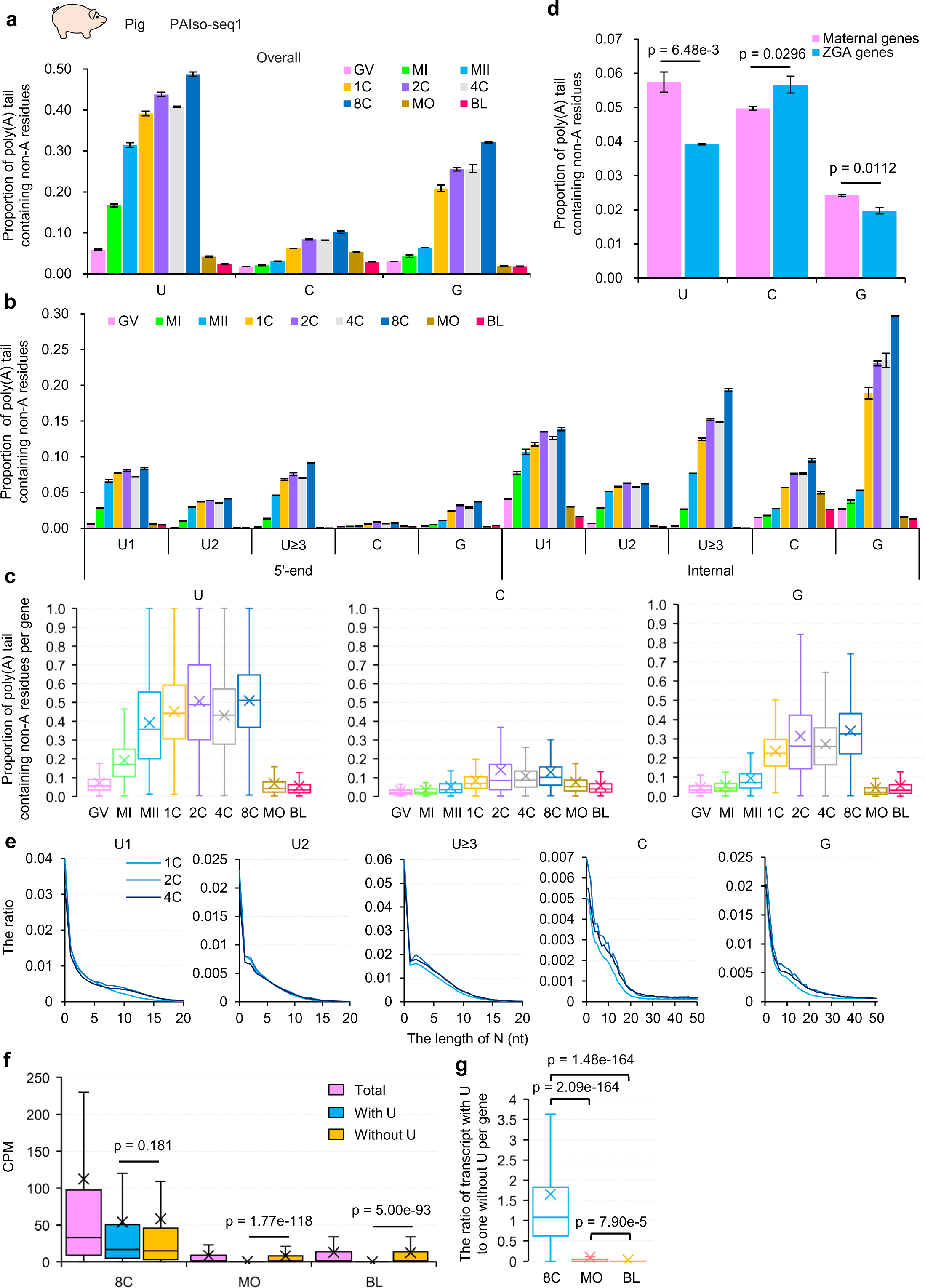
The dynamics of non-A residues during pig OET. **a**, Overall proportion of poly(A) tails containing U, C, and G residues in pig GV, MI, and MII oocytes, and 1C, 2C, 4C, 8C, MO, and BL embryos measured by PAIso-seq1. Transcripts with poly(A) tail of at least 1 nt are included in the analysis. Error bars represent standard error of the mean (SEM) from two replicates. **b**, Proportion of poly(A) tails containing U, C, and G residues in the 5′-end, internal, or 3′-end in different stage pig samples measured by PAIso-seq1. The U residues are further divided according to the length of the longest consecutive U (U1, U2, and U≥ 3). Transcripts with poly(A) tail of at least 1 nt are included in the analysis. Error bars represent SEM from two replicates. **c**, Box plot for the proportion of poly(A) tails containing U, C, and G residues of individual genes in different stage pig samples measured by PAIso-seq1. Genes (n = 3,425) with at least 10 reads and contain poly(A) tails (tail length ≥ 1 nt) in each sample are included in the analysis. **d**, Proportion of poly(A) tails containing U, C, and G residues in maternal and ZGA genes. Error bars represent SEM from two replicates. *P*-value is calculated by Student’s *t* test. **e**, Histogram of the length of N and the ratio of U, C, and G residues in poly(A) tails in pig 1C, 2C, and 4C embryos measured by PAIso-seq1. The U residues are further divided according to the length of the longest consecutive U (U1, U2, and U≥ 3). **f**, Box plot for the expression level of total transcripts, transcripts with U residues, and transcripts without U residues for each maternal gene in pig 8C, MO, and BL embryos measured by PAIso-seq1. Genes (n = 2,929) with at least 10 reads in each sample are included in the analysis. **g**, Box plot for the ratio of number of transcripts with U residues to those without U residues for each maternal gene in pig 8C, MO, and BL embryos measured by PAIso-seq1. Genes (n = 2,929) with at least 10 reads in each sample are included in the analysis. For all box plots, the “×” indicates mean value. The horizontal bars show the median value. The top and bottom of the box represent the value of 25^th^ and 75^th^ percentile, respectively.

**Extended Data Fig. 3.**
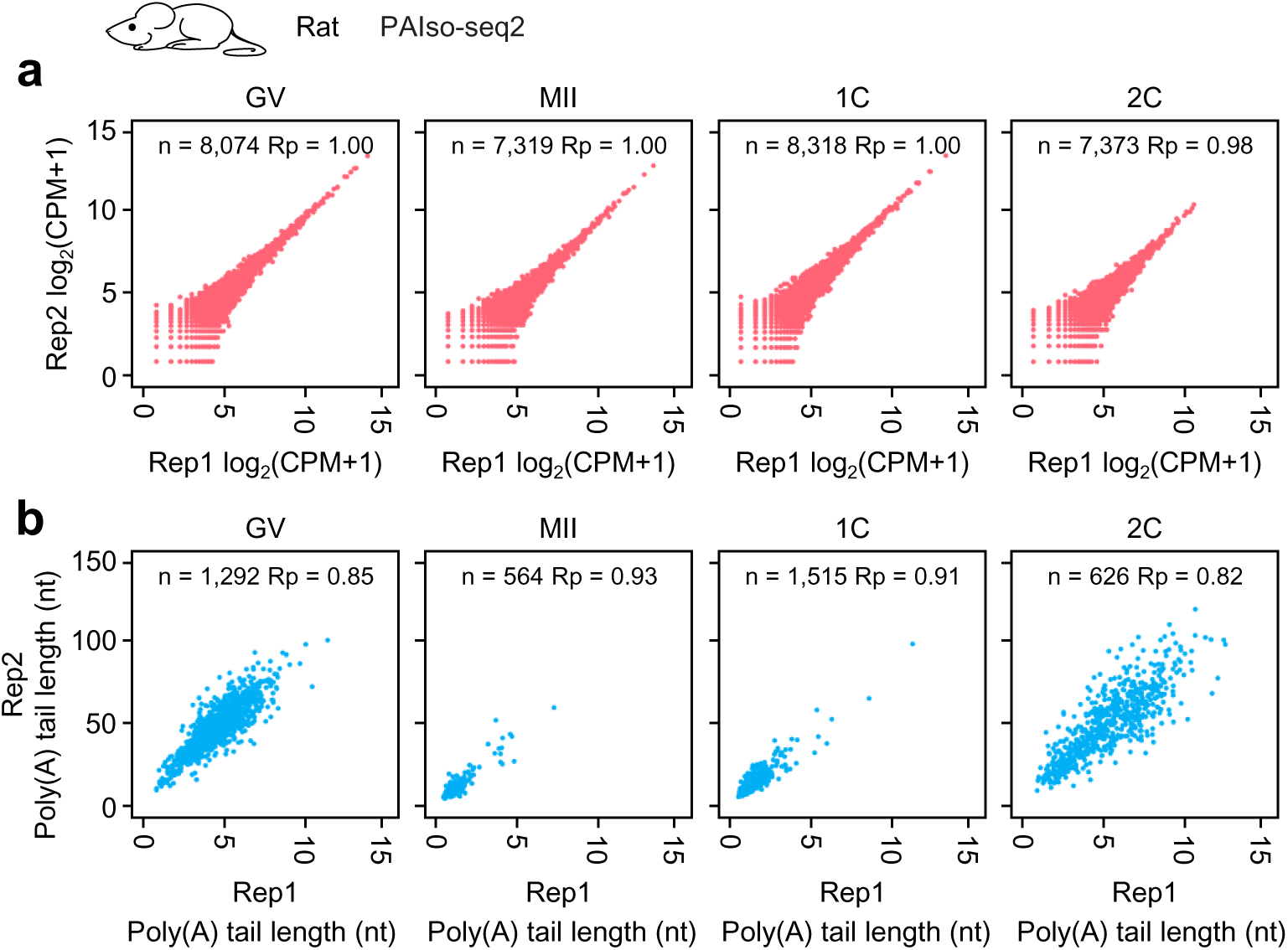
The reproducibility of rat samples. **a**, Scatter plot showing the Pearson correlation of gene expression between two replicates for rat GV and MII oocytes, and 1C and 2C embryos measured by PAIso-seq2. Each dot represents one gene. Pearson’s correlation coefficient (Rp) and the number of genes included in the analysis are shown on the top of each graph. **b**, Scatter plot showing the Pearson correlation of poly(A) tail length between two replicates for rat GV and MII oocytes, and 1C and 2C embryos measured by PAIso-seq2. Each dot represents one gene. The poly(A) tail length for each gene is the geometric mean length of all the transcripts with a poly(A) tail of at least 1 nt for the given gene. Genes with at least 20 reads in each sample are included in the analysis. Pearson’s correlation coefficient (Rp) and the number of genes included in the analysis are shown on the top of each graph.

**Extended Data Fig. 4.**
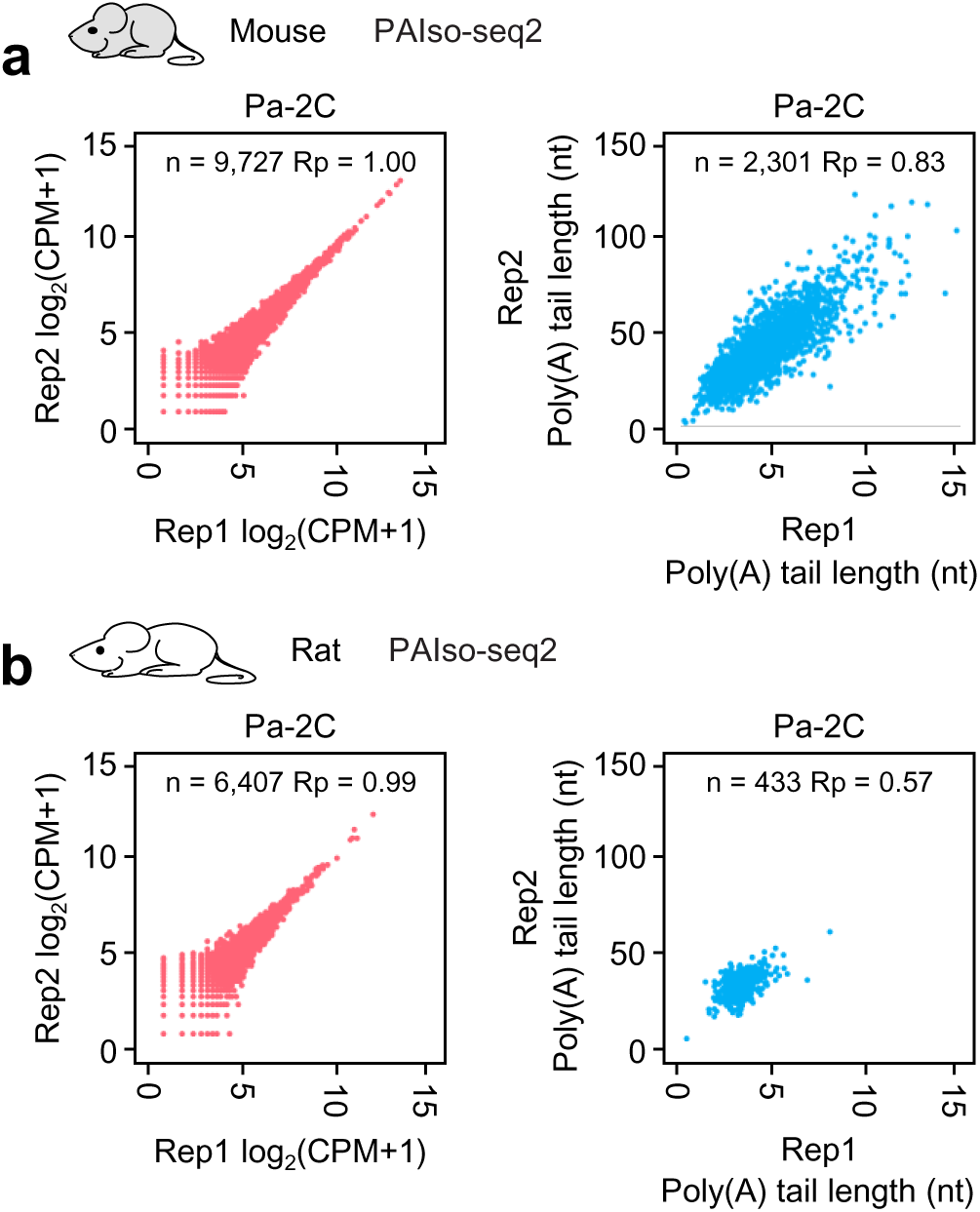
The reproducibility of parthenogenetic samples. Left, Scatter plot showing the Pearson correlation of gene expression between two replicates for 2C and parthenogenetic 2C (Pa-2C) embryos in mouse (**a**) or rat (**b**) measured by PAIso-seq2. Each dot represents one gene. Pearson’s correlation coefficient (Rp) and the number of genes included in the analysis are shown on the top of each graph. Right, Scatter plot showing the Pearson correlation of poly(A) tail length between two replicates for 2C and parthenogenetic 2C (Pa-2C) embryos in mouse (**a**) or rat (**b**) measured by PAIso-seq2. Each dot represents one gene. The poly(A) tail length for each gene is the geometric mean length of all the transcripts with poly(A) tail of at least 1 nt for the given gene. Genes with at least 20 reads in each sample are included in the analysis. Pearson’s correlation coefficient (Rp) and the number of genes included in the analysis are shown on the top of each graph.

**Extended Data Fig. 5.**
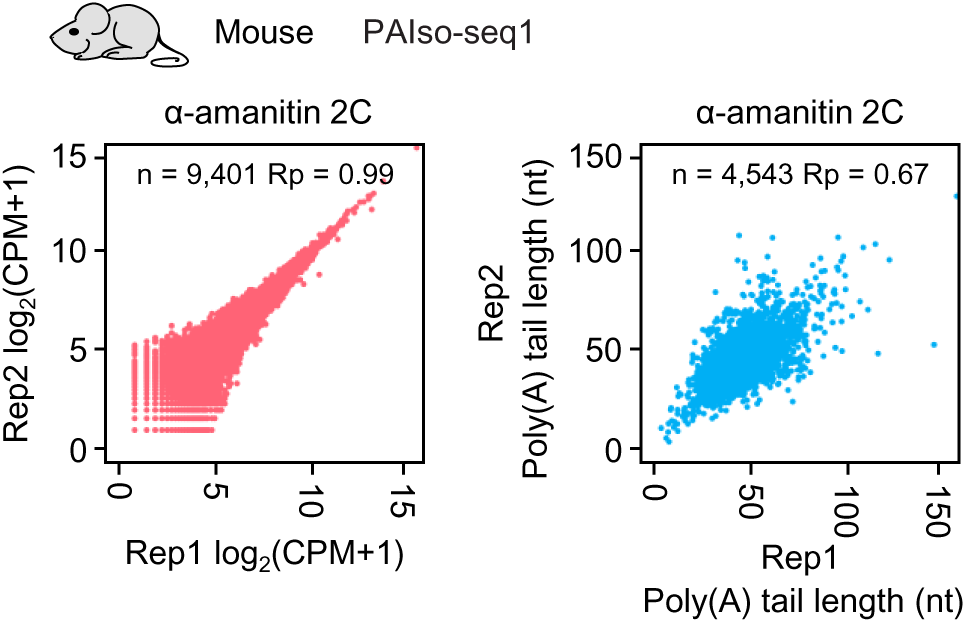
The reproducibility of α-amanitin treated mouse samples. Left, Scatter plot showing the Pearson correlation of gene expression between two replicates for mouse 2C embryos with or without α-amanitin treatment measured by PAIso-seq1. Each dot represents one gene. Pearson’s correlation coefficient (Rp) and the number of genes included in the analysis are shown on the top. Right, Scatter plot showing the Pearson correlation of poly(A) tail length between two replicates for mouse 2C embryos with or without α-amanitin treatment measured by PAIso-seq1. Each dot represents one gene. The poly(A) tail length for each gene is the geometric mean length of all the transcripts with poly(A) tail of at least 1 nt for the given gene. Genes with at least 20 reads in each sample are included in the analysis. Pearson’s correlation coefficient (Rp) and the number of genes included in the analysis are shown on the top of each graph.

